# Genetic structure and differentiation of the endemic Bolle’s Laurel Pigeon (*Columba bollii*) in the Canary Islands

**DOI:** 10.1101/2022.05.31.493538

**Authors:** Patricia Marrero, Rosa Fregel, David S. Richardson

## Abstract

Island archipelagos are global biodiversity hotspots since they often foster high concentrations of diverse lineages and endemic species. Here, we examine the population genetics of the endemic Bolle’s Laurel Pigeon *Columba bollii*, a frugivorous bird inhabiting subtropical laurel forests. We genotyped ten microsatellite loci using DNA obtained from moulted tail feathers collected at eight sampling sites on the four western islands of the Canarian archipelago. Analyses including F-statistics, Bayesian clustering approaches, isolation by distance tests and population graph topologies, were used to infer the genetic diversity and the population differentiation within and among insular populations. Additionally, we evaluated the effect of null alleles on data analysis. Low genetic diversity was found in all populations of Bolle’s Laurel Pigeon, with no significant differences in diversity among them. However, significant genetic differentiation was detected among all populations, with pigeons from La Palma and El Hierro exhibiting the closest affinity. Bayesian clustering supported population separation between islands, and also detected fine-scale structure within the Tenerife and La Gomera populations. Present-day pigeon movements appear to occur between islands, however, this has not been sufficient to remove the signature of genetic divergence among the populations of Bolle’s Laurel Pigeon, which was moderately linked to geographical distance. According to metapopulation structure, this study suggests that the evolutionary history of *C. bollii* is closely related to the geological past of these oceanic islands and the distribution range of its habitat, the laurel forest. Finally, conservation implications for the species are discussed.

## INTRODUCTION

Oceanic islands are exemplary settings for understanding evolutionary and ecological processes (Losos & Ricklefs 2009). Community characteristics of their flora and fauna include disharmony, high endemism and relictualism (Gillespie 2007). Despite the generally discrete geographical status of islands, they are home to a wide diversity of species and habitats (Emerson 2002). Indeed, many insular ecosystems, such as the Macaronesian archipelagos of the Azores, Madeira and the Canaries are global biodiversity hotspots (Myers *et al*. 2000, Cartwright 2019; Florencio *et al*. 2021).

A fundamental source of biodiversity is genetic diversity, which describes the degree of genetic variability within- and between populations (Woodruff 2001). Genetic diversity is a necessary precondition for ecologically relevant traits and provides the means for populations to respond to an ever-changing environment (eg. Hughes *et al*. 2008, Mergeay & Santamaria 2012). Population genetic tools can provide precise estimates of basic features of wildlife populations, such as genetic diversity, gene flow, past historical events and genetic differentiation among populations (Rollins *et al*. 2006, Hohenlohe *et al*. 2021). Microsatellites (also known as simple sequence repeats, SSRs) are codominant genetic markers widely used in population genetic studies as they are highly informative and exhibit high specificity (Vieira *et al*. 2016).

The laurel forest is an endemic subtropical forest of several islands of the Macaronesian region with an outstanding biodiversity richness (del Arco Aguilar *et al*. 2010). These humid forests that covered Eurasia and the Mediterranean Basin from the early Cenozoic until the end of the Pliocene, show refuge and stepping-stone dispersal (Fernández-Palacios *et al*. 2011, Bogaard 2013). Given the diverse biogeographical events occurring over this prolonged geological timescale, insular laurel forests are today composed of plant species with a heterogeneous origin and high taxonomic uniqueness (Médail & Quézel 1999, Fernández-Palacios *et al*. 2011, Kondraskov *et al*. 2015). Species with fleshy fruits, belonging to the genera *Apollonias, Laurus, Ocotea, Persea* (Lauraceae), *Picconia* (Oleaceae), *Prunus* (Rosaceae) and *Visnea* (Theaceae) still co-occur in the laurel forests. The recurrent long-distance dispersal of these plant species to Macaronesia and the establishment of some plant lineages would have been facilitated by endozoochory (Vargas 2007). By ingesting and excreting and/or regurgitating viable seeds, fruit-eating animals (mainly birds) are able to promote the germination and disperse the seeds of these fleshy fruiting plants (Hampe 2003).

The endemic Bolle’s Laurel Pigeon (*Columba bollii*) is part of this frugivore guild in the laurel forests on the western islands (Tenerife, La Gomera, La Palma and El Hierro) of the Canary archipelago. Although the eastern islands (Gran Canaria, Fuerteventura and Lanzarote) harboured laurel forests in the past, only now trace relicts remain due to human activities (Parsons 1981, Martín Osorio *et al*. 2011). The few Columbidae fossils that have been found on these eastern islands have not been specifically identified (Alcover & Florit 1989, Rando & Perera 1994). Currently, *C. bollii* shares habitat and insular distribution with the also endemic White-tailed Laurel Pigeon *C. junoniae*, however it is phylogenetically closer to the Madeira Laurel Pigeon *C. trocaz*; both deriving from a common ancestor with the European Woodpigeon *C. palumbus* (Dourado *et al*. 2014).

The close ecological linkage between *C. bollii* and the laurel forests is supported by the spatio-temporal fluctuations recorded in diet and local abundance of pigeons, and the differential availability of fruit resources (Martín *et al*. 2000, Marrero 2009, Marrero & Nogales 2021). Pigeons can move among forest patches in response to their fruiting phenology and food preferences (Martín *et al*. 2000). However, we still lack detailed information on the dispersal of *C. bollii* and how this may affect connectivity among its populations. Such data would ultimately determine the degree of gene-flow cohesion with the structure of genetic diversity. To examine the genetic consequences of distribution of pigeon populations in the fragmented laurel forest, population genetic analysis is essential.

Here we used microsatellite markers applied to DNA derived from non-invasive samples to investigate the population genetic structure of the elusive and poorly studied *C. bollii*. In this case, moulted feathers collected in the field minimize the stress and disturbance to the birds and provide an appropriate source of DNA (Presti *et al*. 2013). The goals of this study were to: (i) determine genetic structure among populations of *C. bollii* within and among islands, (ii) assess patterns of diversity and genetic differentiation between populations, (iii) evaluate rates of gene flow and therefore, effective dispersal among populations, and (iv) discuss whether the genetic population data are indicators of the evolutionary relationship between this endemic pigeon and its ancestral habitat.

## METHODS

### Study area and sample collection

Fieldwork was conducted in 2007-2008 and 2011 across all laurel forest remnants within the distribution range of *C. bollii*. These ancient forests occupy only 18% (16,419 Ha) of their original area in the Canary archipelago, due to progressive human-mediated deforestation (Del Arco Aguilar *et al*. 2010, Fernández-Palacios *et al*. 2011). Although the laurel forest has been reduced, modified or otherwise almost extirpated on all the islands, fragmentation is clearly evident on Tenerife, where three main relict forests can be identified: Anaga (T-AN), in the north-east of the island, La Esperanza-Tigaiga (T-ET) in the north, and Teno (T-TE) in the north-west (Fig. 1A). For the sampling of La Gomera and La Palma, we considered that the deep ravine areas in the north of these islands could act as barriers. We therefore sampled across two potentially divided areas on each island: Hoya del Tión-Los Roques (G-HR) in the north and Epina-Los Pajaritos (G-EP) in the south of La Gomera, and Barlovento-Niquiomo (P-BN) in the east and Garafía-Tinizara (P-GT) in the north-west of La Palma. Samples collected on El Hierro were treated as from a single population, Jinama-Tina de las Casillas (H-JT), since most of the laurel forest lies on a slope with a smoother relief (Fig. 1A). Across these eight sampling sites, moulted feathers of pigeons were collected and stored in individual paper envelopes or plastic bags under dark dry conditions. Feathers from birds killed by predators were also opportunistically sampled. In all cases, only fresh samples were analysed. Finally, GPS coordinates for each sample were recorded.

**Figure 1.**
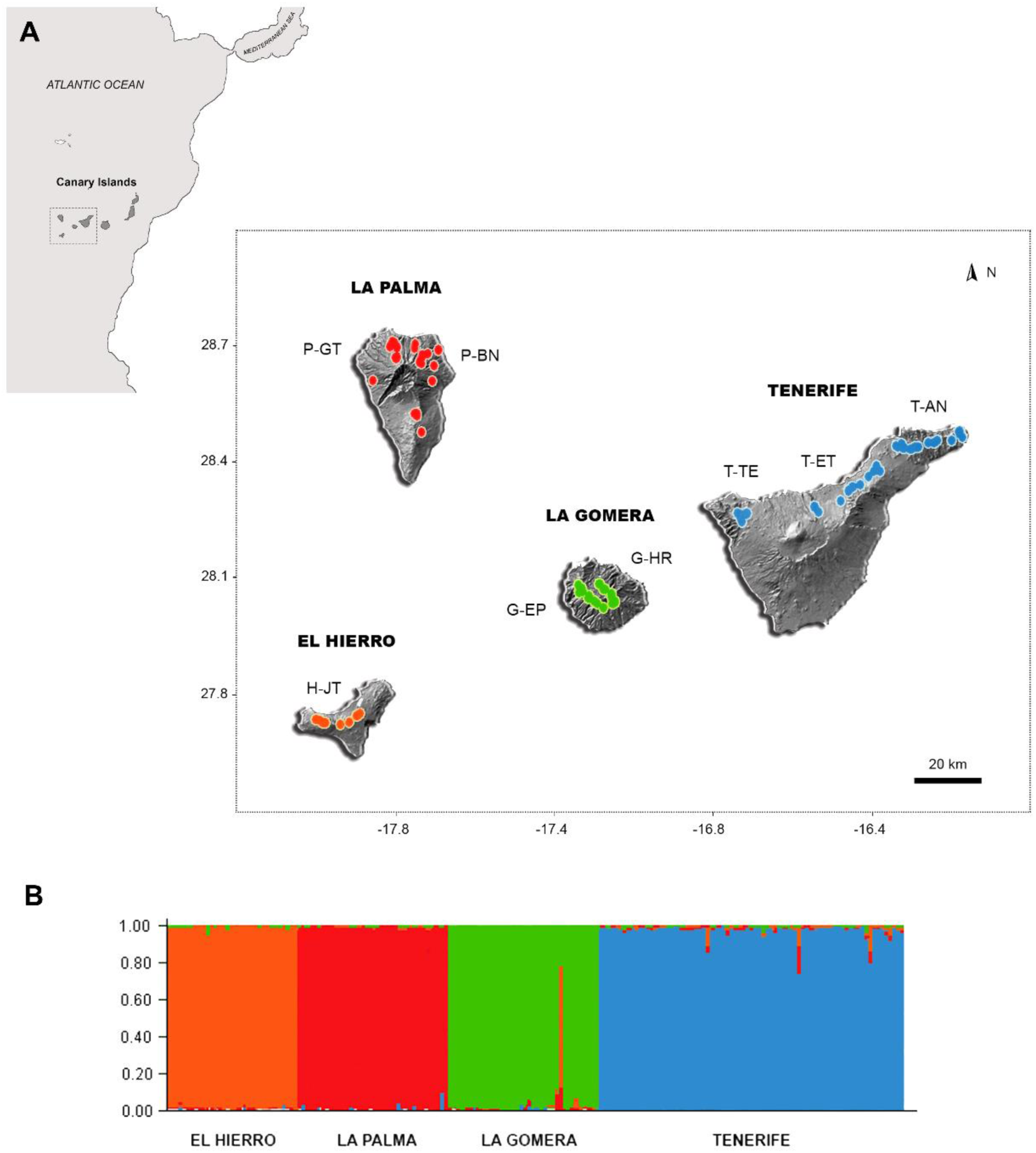
Map of the Canary archipelago and assigned genetic clusters of Bolle’s Laurel pigeon (*C. bollii)* from STRUCTURE. (A) Sampling localities in El Hierro (H-JT: Jinama-Tina de las Casillas), La Palma (P-BN: Barlovento-Niquiomo and P-GT: Garafía-Tinizara), La Gomera (G-EP: Epina-Los Pajaritos and G-HR: Hoya del Tión-Los Roques), and Tenerife (T-AN: Anaga, T-ET: La Esperanza-Tigaiga and T-TE: Teno). (B) Results of the Bayesian analysis by admixture model showing four clusters as the highest Delta k value for Bolle’s Laurel pigeon populations. Each bar shows a single individual, with each colour representing the proportion of a given genetic cluster to an individual’s genotype.

### DNA extraction and sexing test

Individual feathers were chosen for genomic DNA extraction based on two criteria. First, large feathers were selected as they provide more DNA than smaller feathers (Bayard de Volo *et al*. 2008). Secondly, tail feathers were chosen as this allowed us to accurately resolve between *C. bollii* (whose feathers are blackish with a broad pale-grey subterminal band) and the sympatric *C. junoniae* (brownish tail feathers with a creamy-white terminal band). This visual diagnostic character avoids the additional expense associated with prior species identification of samples (Marrero *et al*. 2008). DNA extraction was performed using the DNeasy Blood & Tissue kit (Qiagen), following the manufacturer’s instructions with slight modifications. All surfaces and laboratory supplies were routinely cleaned with 10% bleach and then rinsed in sterile double-distilled water to avoid contamination among samples (Eggert *et al*. 2005). The entire calamus of each tail feather, including the blood clot of the superior umbilicus (Horváth *et al*. 2005), was cut transversely into small pieces and incubated with agitation overnight at 56ºC in 180-200 µl of ATL buffer (lysis solution) and 20 µl of proteinase K (20 mg/ml). DNA concentration and purity were determined for each extract using a Nanodrop spectrophotometer (NanoDrop 8000, Thermo Scientific, Wilmington, DE). PCR was performed with primers 18S-F/18S-R (Wu *et al*. 2007), to check whether DNA extracts were amplified satisfactorily. This primer pair generates a 256-base-pair (bp) fragment from the 18S ribosomal gene in both male and female Columbidae, which also served as internal control for sex determination (Wu *et al*. 2007). Amplifications were carried out in 10 µl total volume, containing 10-15 ng of template DNA, 0.5 µM of each primer, 10x CoralLoad PCR buffer and 5 µl of TopTaq Master Mix 2x (Qiagen). An initial denaturation cycle of 94°C for 6 min was followed by 35 cycles of 94°C for 45 s, 58°C for 45 s and 72°C for 45 s. All reactions ended with a final extension cycle at 72°C for 10 min. PCR products were visualised on 2% agarose gels stained with ethidium bromide.

The conserved primers P8/P2 based on the CHD-Z and CHD-W genes (Griffiths *et al*. 1998) were used to determine sex. The reaction mixture was similar to those of 18S-F/18S-R, but including 20 mg/ml BSA and additional MgCl^2^ (2.0 mM). PCR conditions were 94ºC for 3 min, 40 cycles of 94 ºC for 30 s, 55 ºC for 45 s and 72 ºC for 45 s, followed by 72 ºC for 10 min and a final step of 20 ºC for 60 s. PCR products were resolved on a 4% agarose gel stained with ethidium bromide. Negative controls were included in all amplification runs. Due to the lack of preserved specimens of both sexes of *C. bollii*, we used muscle tissue from carcasses of *C. trocaz* (its closest relative), sexed by gonadal analysis, as reference positive control. Two PCR repetitions were performed when was necessary to ensure the correct sex assignment of each feather sample.

### Microsatellite markers and genotyping

Since no information on the genomic sequence of *C. bollii* is available, we used the conserved microsatellite markers developed by Dawson *et al*. (2010) that can be utilised across a wide range of bird species. All single-plex PCR reactions were performed in a final volume of 2 μl with 0.2 µM of each primer, 1 µl Qiagen Multiplex PCR Master Mix, and approximately 12 ng of template DNA. The general PCR conditions were 15 min at 95°C, 40 cycles of 30 s at 94°C, 90 s at 58°C and 60 s at 72°C, followed by a final extension step at 60°C for 30 min. Products were diluted (1/200) and analysed on an ABI 3730 capillary sequencer (Applied Biosystems). Alleles were scored using GENEMARKER 2.4.0 (SoftGenetics) and GENEMAPPER 4.0 (Applied Byosystems). In order to detect contamination, a negative control was included in the amplifications. For some samples, re-extraction, ethanol precipitation and/or dilution of template DNA were also required to solve PCR inhibition problems.

### Population genetics analyses

The reliability of genotyping results was estimated using a rarefaction method implemented in the R package “vegan” (Oksanen *et al*. 2013), to correct sample size differences. To assess genetic diversity within populations, the mean number of alleles per locus (A), number of private alleles (Ap), and observed (H_O_) and unbiased expected heterozygosities (H_E_) for each locus and population were calculated using GENALEX 6.5 (Peakall & Smouse 2012). The software FSTAT 2.9.3.2 (Goudet 2002) was used to calculate allelic richness (Ar) and fixation index (F_IS_). Deviations from Hardy-Weinberg (HWE) and linkage equilibrium for each locus and population were estimated in GENEPOP 4.0 (Raymond & Rousset 1995), using the exact test with default values for the Markov chain parameters (1,000 dememorisation steps, 100 batches and 1,000 iterations per batch). A sequential Bonferroni correction for multiple testing was applied to adjust p-values (Rice 1989). The microsatellite genotyping from non-invasive samples containing low quantities of amplifiable DNA can result in heterozygote deficits caused by the presence of null alleles (Broquet & Petit 2004). Thus, the frequency of null alleles 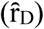 for each locus was estimated using the EM algorithm of Dempster *et al*. (1977) with 10,000 bootstrap repetitions implemented in the program FREENA (Chapuis & Estoup 2007). To check the effects of null alleles on our genetic diversity results, genotypes were corrected according to the estimated false homozygotes, and H_O_, H_E_, F_IS_ statistics and HWE test recalculated (Chapuis & Estoup 2007, Oddou-Muratorio *et al*. 2009, Sun *et al*. 2012).

Pairwise F_ST_ and D_est_ estimators quantified genetic divergence among populations over loci. The measure of genetic fixation, F_ST_, is calculated as the ratio of the variance in allele frequencies among populations to the overall variance (Weir & Cockerham 1984). Additionally, D_est_ measures genetic differentiation among populations corrected for sampling bias using allele-diversity (Jost 2008), which is thought to be more appropriate for microsatellite data sets (Bird *et al*. 2011). F_ST_ values were estimated in GENALEX 6.5 (Peakall & Smouse 2012) while D_est_ was calculated in the “DEMEtics” package for R (Gerlach *et al*. 2010). Statistical significance and 95% confidence intervals were tested by 10,000 bootstrap repetitions. FREENA was also used to obtain pairwise F_ST_ corrected for null alleles, F_ST_^{ENA}^ (Chapuis & Estoup 2007). Hierarchical analyses of molecular variance (AMOVA) with 10,000 permutations were calculated using ARLEQUIN 3.5.1.3 (Excoffier & Lischer 2010) to assess genetic variance at two levels: among islands, and among populations within islands. Based on the Cavalli-Sforza and Edwards distances (D_c_), genetic relatedness among samples was depicted by an unrooted Neighbour-Joining (NJ) tree using POPULATION 1.2.32 (Langella 2002), and visualised in FIGTREE 1.4.2 (Rambaut 2007). A Principal Component Analysis (PCA) of the microsatellite individual-genotype matrix was performed using PCAGEN (Goudet 1999) to identify spatial patterns of genetic variation.

Population structure was determined using a Bayesian algorithm-based method in STRUCTURE 2.3.4 (Pritchard *et al*. 2000, 2010). The prior population information model was used, considering each island as a defined population. However, the admixture model was also applied at an initial point of the analysis, as recommended by the authors (Pritchard *et al*. 2010). Ten independent runs of K (number of clusters) for 1-10 possible populations were performed, using correlated allele frequencies between populations. Burn-in period and Markov Chain Monte Carlo (MCMC) repetitions were 50,000 and 300,000 respectively. STRUCTURE HARVESTER 0.6.93 (Earl & vonHoldt 2012) was used to establish the value of K that best fits the data set deriving from the Evanno method (Evanno *et al*. 2005). First generation migrants were estimated in GENECLASS2 (Piry *et al*. 2004). The likelihood that an individual belongs to the reference population was calculated using the ratio of the population where the individual was sampled (L_home) to the highest likelihood observed in all sampled populations (L_max), described by Paetkau *et al*. (2004). The computation method was based on frequencies (Paetkau *et al*. 1995) and the Monte Carlo resampling algorithm was run on 10,000 simulated individuals, taking into account all loci and a type I error of 0.01.

A Mantel test was performed on log-transformed genetic and geographical distances in IBDWS 3.23 with 10,000 permutations (Jensen *et al*. 2005). The chord distance (D_C_) of Cavalli-Sforza & Edwards (1967), calculated by the INA method in FREENA, was used as a measure of genetic distance between populations, due to its limited sensitivity to the presence of null alleles (Chapuis & Estoup 2007). Geographical distances were calculated (in km) from UTM coordinates using Geographic Distance Matrix Generator 1.2.3 (Ersts 2013).

Finally, Population Graph analysis was also used to explain genetic population structure. It is a multivariate graph-theoretical approach in which nodes (populations) are connected in a network by the shortest path of the edges, named conditional genetic distance (cGD). The edge width represents the degree of dependence of evolutionary trajectories between populations, while node sizes indicate allelic richness within populations (Dyer & Nason 2004). The significance level was adjusted for *P* = 0.05. This method was implemented in R package “popgraph” (Dyer, 2013). In a simulation analysis, Klütsch *et al*. (2012) found that null alleles influence population graph topologies, independently of their frequency in the populations. For that reason, the null allele corrected data set was also evaluated. Evidence of isolation by distance was also examined using the genetic covariance.

## RESULTS

### DNA extraction and sexing

DNA was extracted from 186 field-sampled moulted tail feathers: Tenerife (77; T-AN: 37, T-ET: 19 and T-TE: 21), La Gomera (38; G-EP: 23 and G-HR: 15), La Palma (38; P-BN: 21 and P-GT: 17) and El Hierro (H-JT: 33). The integrity of DNA obtained from feathers was demonstrated by the 100% amplification success rate for the 18S ribosomal DNA. Molecular sexing reactions provided a reliable DNA profile allowing for gender identification of *C. bollii* (Fig. 2): Amplicons from females consisted of two different sized bands (∼ 350 and ∼ 370 bp), while males showed only the larger band (∼ 370 bp). We were able to sex 65.6% of total samples, of which 54.1% were identified as females and 45.9% as males.

**Figure 2.**
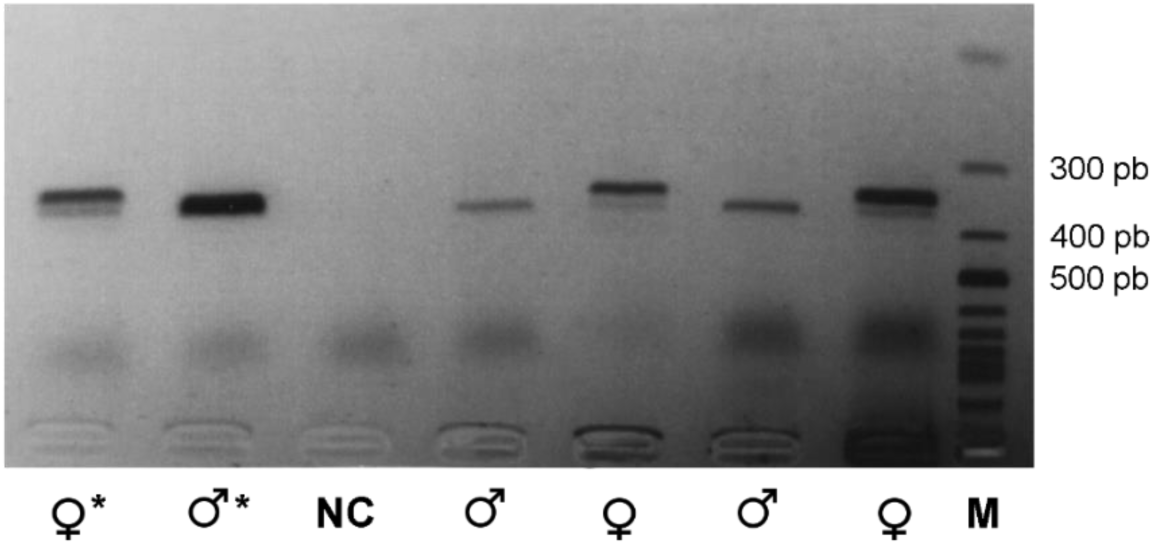
Feather sexing test of Bolle’s Laurel pigeon (*C. bollii)* using PCR products amplified for CHD-Z and CHD-W genes by primer sets P2/P8. Females (♀) have two bands and males (♂) one band. M is a 1Kb DNA marker, NC the negative control and the asterisk indicates the positive controls.

### Microsatellite analysis and genetic diversity

Of the 34 markers tested, 10 yielded polymorphic products within *C. bollii* (TG01-077, TG01-124, TG01-148, TG02-078, TG02-120, TG03-002, TG04-012, TG07-022, TG13-017 and TG22-001), with allelic ranges similar to that described by Dawson *et al*. (2010) for other avian species. The rarefaction analysis demonstrated that sample sizes were clearly adequate to detect the cumulative allelic richness of each population (Fig. S1).

A total of 60 alleles were identified across 10 microsatellite loci (Table 1), with a mean per locus of 3.85 and an allelic richness of 3.57 for all islands, estimated from 23 diploid individuals as a minimum sample size. The maximum and minimum numbers of unique alleles were found on Tenerife (10) and El Hierro (3), respectively. The average observed and unbiased expected heterozygosities were low, ranging from 0.233 to 0.270 and 0.325 to 0.374, respectively. The deficiency in the proportion of observed heterozygotes with respect to the expected is also reflected by positive fixation indices (F_IS_ for all islands = 0.272, 95% CI = 0.195-0.358). Similar results were obtained at population level, except for the number of private alleles, which was higher in G-HR (7) and absent in P-BN and G-EP (Table 1). No significant difference was found in the genetic diversity measures (Ar, H_O_, H_E_ and F_IS_) between islands (*P* > 0.05 for paired t-test) or populations. Genetic diversity values per locus are shown in Table S1. Significant deviations from HWE after Bonferroni correction were detected in some loci but were not equally represented on all islands (Table S1). On Tenerife, six loci were out of HWE, while for La Palma, El Hierro and La Gomera only one to four were not at equilibrium. At population level, most of the deviation derived from T-AN samples on Tenerife, with five loci in disequilibrium. The departure from HWE was not locus specific. A monomorphic locus (TG07-022) was also detected for La Palma. There was no significant pattern of linkage disequilibrium (*P* > 0.05) for pairs of loci.

**Table 1.** Summary statistics of genetic diversity derived using ten microsatellite loci for *C. bollii* populations in the Canary Islands. N = number of samples, *A* = mean number of alleles per locus, *Ar* = mean allelic richness, *Ap* = number of private alleles, *H*_*O*_ = observed heterozygosity, *H*_*E*_ = unbiased expected heterozygosity and *F*_*IS*_ = fixation index. ^C^ Statistics corrected for null alleles. Sampling localities are indicated as H-JT: Jinama-Tina de las Casillas in El Hierro, P-BN: Barlovento-Niquiomo and P-GT: Garafía-Tinizara in La Palma, G-EP: Epina-Los Pajaritos and G-HR: Hoya del Tión-Los Roques in La Gomera, and T-AN: Anaga, T-ET: La Esperanza-Tigaiga and T-TE: Teno in Tenerife.

| Island/Population | *N* | *A* | *Ar* | *Ap* | *H<sub>O</sub>*<sup>c</sup>/*H<sub>O</sub>* | *H<sub>E</sub>*<sup>c</sup>/*H<sub>E</sub>* | *F<sub>IS</sub>*<sup>c</sup>/*F<sub>IS</sub>* |
| --- | --- | --- | --- | --- | --- | --- | --- |
| **El Hierro** | **33** | **3.70** | **3.60** | **0.30** | **0.255/0.436** | **0.374/0.429** | **0.321/-0.015** |
| H-JT | 33 | 3.70 | 3.12 | 0.30 | 0.255/0.436 | 0.374/0.429 | 0.321/-0.015 |
| **La Palma** | **38** | **3.90** | **3.55** | **0.40** | **0.270/0.398** | **0.342/0.403** | **0.214/0.012** |
| P-BN | 21 | 3.10 | 2.79 | 0.00 | 0.268/0.376 | 0.324/0.369 | 0.178/-0.020 |
| P-GT | 17 | 3.50 | 3.25 | 0.30 | 0.271/0.424 | 0.361/0.427 | 0.254/0.008 |
| **La Gomera** | **38** | **3.60** | **3.20** | **0.70** | **0.233/0.396** | **0.325/0.407** | **0.286/0.029** |
| G-EP | 23 | 3.00 | 2.71 | 0.00 | 0.211/0.379 | 0.307/0.383 | 0.317/0.009 |
| G-HR | 15 | 3.20 | 3.11 | 0.70 | 0.276/0.388 | 0.357/0.400 | 0.234/0.031 |
| **Tenerife** | **77** | **4.20** | **3.29** | **1.00** | **0.249/0.441** | **0.340/0.441** | **0.269/0.001** |
| T-AN | 37 | 3.60 | 2.86 | 0.30 | 0.205/0.467 | 0.347/0.470 | 0.419/0.007 |
| T-ET | 19 | 2.80 | 2.62 | 0.10 | 0.298/0.411 | 0.325/0.382 | 0.084/-0.080 |
| T-TE | 21 | 3.20 | 2.94 | 0.30 | 0.285/0.393 | 0.343/0.398 | 0.174/0.014 |
| **All islands** | **186** | **3.85** | **3.57** | **0.60** | **0.252/0.418** | **0.345/0.420** | **0.272/0.007** |

The frequency of null alleles per locus per island ranged from 0 - 0.291, with a mean frequency of 0.084. TG02-078 showed the highest mean frequency of null alleles 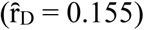 and TG03-002 the lowest 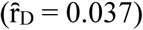, although these values varied according to island (Table S1). The correction of the data set for null alleles increased heterozygosity values (H_O_ = 0.396-0.436 and H_E_ = 0.403-0.429), reduced the overall heterozygote deficit (F_IS_ = 0.007, 95% CI -0.010-0.025), and all islands/populations and loci were in Hardy-Weinberg equilibrium.

### Genetic population differentiation

Genetic divergence, estimated as pairwise F_ST_ and D_est_ between islands, ranged from 0.008 to 0.020 for F_ST_, and from 0.0008 to 0.018 for D_est_. All pairwise comparisons were significant after sequential Bonferroni correction, except those involving samples from La Palma and El Hierro (Table 2). Both estimators gave similar results, however, pairwise F_ST_^{ENA}^ suggests that null alleles underestimated the overall differentiation between islands (Table 3). When the impact of null alleles was estimated on pairs of populations, the genetic divergence was more evident (Table S2). Highly significant differences (P<0.01) were found between La Gomera populations (G-EP and G-HR) and those from El Hierro (H-JT) and La Palma (P-BN and P-GT), while T-AN and T-ET were generally different from the other insular populations. The global F_ST_ was 0.015 (*P* = 0.001, 95% CI 0.006-0.025) but the overall differentiation excluding null alleles was slightly higher with F_ST_^{ENA}^ = 0.024 (*P* < 0.001, 95% CI 0.011-0.039).

**Table 2.** Pairwise *F*_*ST*_ (below diagonal) and *D*_*est*_ values (above diagonal) between the four genetic clusters of *C. bollii* in the Canary Islands. 95% confidence intervals are shown.

|  | El Hierro | La Palma | La Gomera | Tenerife |
| --- | --- | --- | --- | --- |
| El Hierro |  | 0.0008 (-0.006-0.014)<sup>n.s.</sup> | 0.015 (0.008-0.028)* | 0.018 (0.014-0.026)* |
| La Palma | 0.008<sup>n.s.</sup> |  | 0.013 (0.007-0.022)* | 0,011 (0.007-0.019)* |
| La Gomera | 0.020* | 0.019* |  | 0.010 (0.006-0.017)* |
| Tenerife | 0.018** | 0.013* | 0.014* |  |

\* $P < 0.05$ , \*\* $P < 0.01$ , n.s. = not significant

**Table 3.** Pairwise *F*_*ST*_^*{ENA}*^ (below diagonal) and their significance (above diagonal) between the populations of *C. bollii* on the Canary islands. 95% confidence intervals are shown.

|  | El Hierro | La Palma | La Gomera | Tenerife |
| --- | --- | --- | --- | --- |
| El Hierro |  | n.s. | *** | *** |
| La Palma | 0.002 (-0.003-0.009) |  | *** | *** |
| La Gomera | 0.046 (0.008-0.103) | 0.039 (0.009-0.080) |  | *** |
| Tenerife | 0.025 (0.013-0.038) | 0.019 (0.006-0.032) | 0.019 (0.003-0.036) |  |

\*\*\* $P < 0.001$ , n.s. = not significant

AMOVA results as a weighted average over loci revealed that 1.64% (F_CT_ = 0.016, *P* = 0.001) of the total genetic variation was attributable to the variability among islands, 0.39% (F_SC_ = 0.003, *P* = 0.613) to the variability between populations within islands and 97.98% (F_ST_ = 0.020, *P* = 0.004) was accumulated within populations within islands. Clustering analysis using the NJ algorithm from D_c_ genetic distances showed that samples from La Gomera and Tenerife clustered together in their respective branches, while those from La Palma were mainly associated with El Hierro (Fig. 3A). The first two PCA axes performed on the allele frequencies of eight populations explained about 60% of the genetic variability among populations. This also implied that populations from geographically close islands did not cluster together, although populations from La Palma were closer to the El Hierro population on the PC1-axis (Fig. 3B).

**Figure 3.**
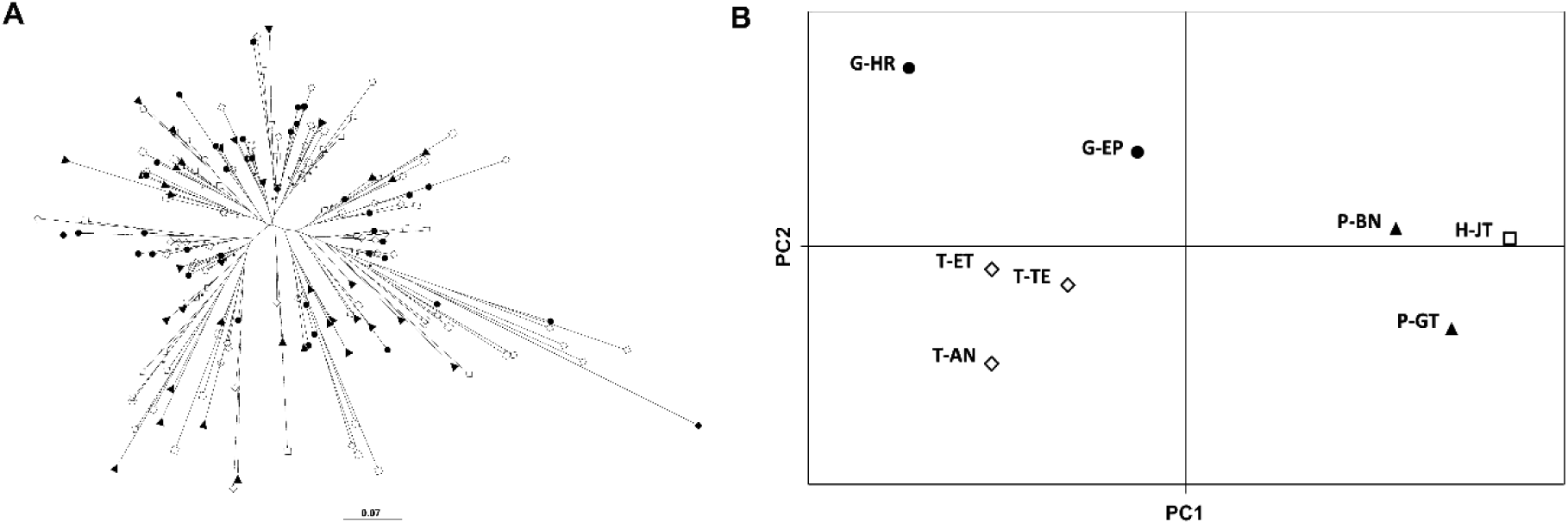
Clustering of Bolle’s Laurel pigeon (*C. bollii*) based on Neighbor Joining (NJ) and Principal Component Analysis (PCA). (A) Unrooted NJ tree inferred from Cavalli-Sforza genetic distances. Scale bar represents branch lengths. (B) PCA plot of averaged first (PC1) and second (PC2) principal component scores for the eight pigeon populations. Island distribution is shown with different colours and symbols (white square: El Hierro, black triangle: La Palma, black circle: La Gomera, and white diamond: Tenerife). Sampling localities are indicated as H-JT: Jinama-Tina de las Casillas in El Hierro, P-BN: Barlovento-Niquiomo and P-GT: Garafía-Tinizara in La Palma, G-EP: Epina-Los Pajaritos and G-HR: Hoya del Tión-Los Roques in La Gomera, and T-AN: Anaga, T-ET: La Esperanza-Tigaiga and T-TE: Teno in Tenerife.

The two models run in STRUCTURE 2.3.4 were largely consistent in the number of genetic clusters (K) estimated. The admixture model gave K = 4 as the most probable number of clusters and secondary peaks with larger standard deviations at K = 3, 6 and 9. Similarly, K = 4 was clearly supported by the prior population information model, when four islands were assumed (Fig. 1B). The genetic structure analysis within each cluster showed support for K = 1 for La Palma and El Hierro, K = 2 for La Gomera and K = 3 for Tenerife (Figure S2).

To evaluate the effect of null alleles on the STRUCTURE analysis, TG02-078 (with the highest frequency of null alleles) was deleted from the genotype data set and all the analyses rerun. The overall data then showed that the optimal number of clusters was three using the admixture model, but four using the prior population information model. The same pattern of sub-clusters within islands was obtained (results not shown). GENECLASS2 test was consistent with the STRUCTURE results because most individuals were assigned to the population in which they were sampled. Nevertheless, four possible first-generation migrants were detected; two of them from La Palma to El Hierro and La Gomera, and two from Tenerife to La Gomera and La Palma. No evidence of recent movements within islands was detected, possibly due to low power as a result of small sample size. However, these results mainly reflect affinities among individuals, rather than true migration events. The Mantel test of isolation by distance showed a positive relationship between geographical and genetic distances using D_C_ (*r* = 0.182, *P* = 0.178) and cGD (*r* = 0.432, *P* = 0.004) for the eight populations.

The Population Graph analysis showed 14 edges connecting the eight populations (*p*_*edge*_ = 0.5), which represent the best-fit model to the total among-population covariance structure (Fig. 4). The lower the number of edges, the higher the conditional genetic distance between nodes. The highest degree of connectivity was shown by T-TE, while T-AN was the most isolated population. Network topology also suggests that spatial genetic structure may depend on other variables besides geographic distances among populations, since populations geographically distant from each other do not always present higher genetic differentiation. This is shown by the thickness of the edges in Fig. 4. Analysis using the data set corrected for null alleles resulted in a different network structure (Fig. S3): a lower number of edges (*p*_*edge*_ = 0.393) reflected the separation of La Palma and El Hierro populations from those of La Gomera, with conditional independence of Tenerife populations through the links of P-BN and P-GT with T-AN and T-TE, respectively. In any case, the Population Graph revealed that La Palma and El Hierro populations constituted a module or complete graph subset (Fig. S3).

**Figure 4.**
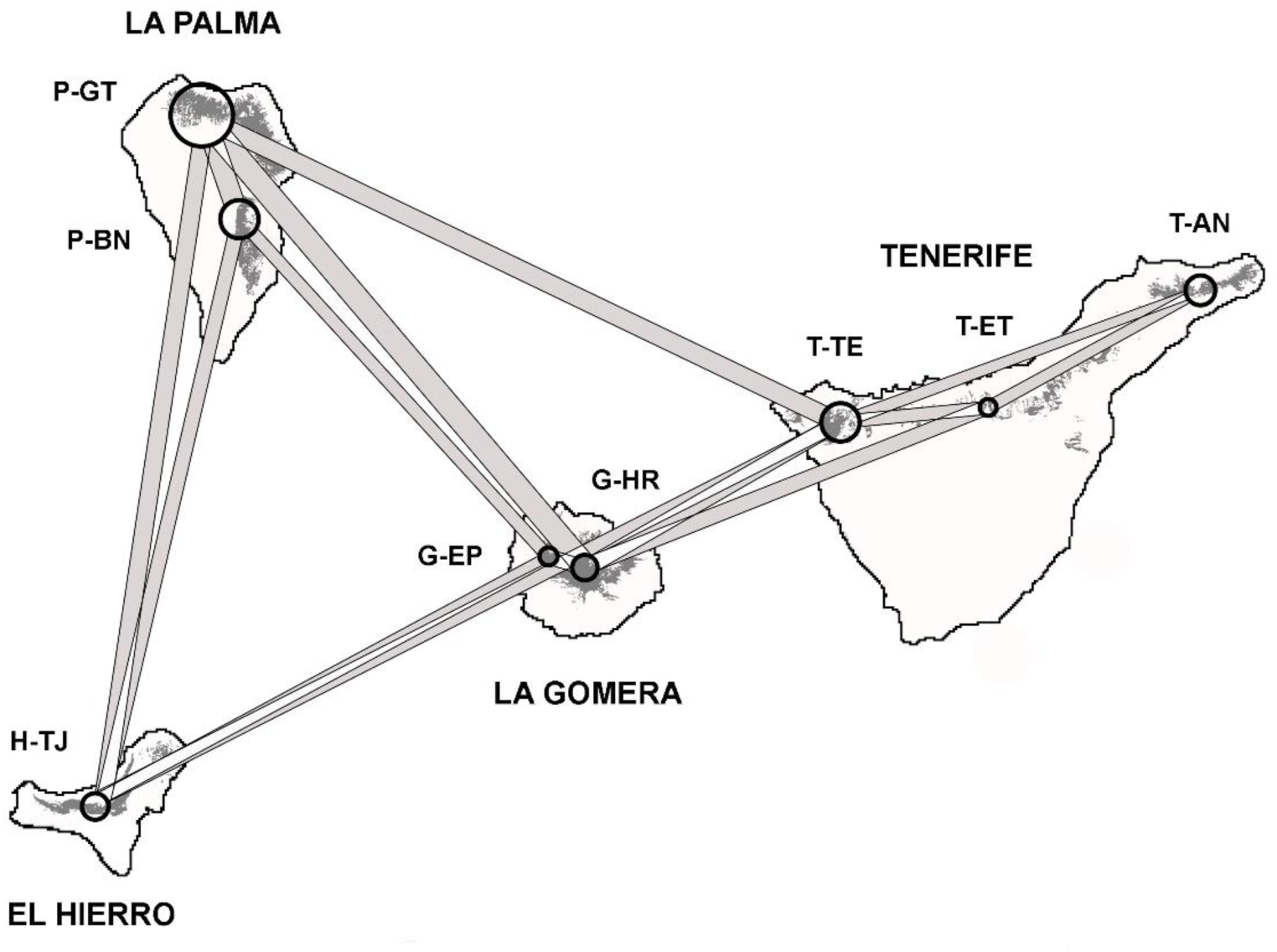
Population graph showing the connectivity network among Bolle’s Laurel pigeon (*C. bollii)* populations within laurel forest on the Canary Islands. Circles represent the central location of sampled populations (size is proportional to within-population allelic richness). The grey edge set indicates the genetic structure conditionally dependent among populations (line width shows the strength of genetic covariance). Dark grey areas represent the current areas of laurel forests (Del Arco *et al*. 2010). Sampling localities are indicated as H-JT: Jinama-Tina de las Casillas in El Hierro, P-BN: Barlovento-Niquiomo and P-GT: Garafía-Tinizara in La Palma, G-EP: Epina-Los Pajaritos and G-HR: Hoya del Tión-Los Roques in La Gomera, and T-AN: Anaga, T-ET: La Esperanza-Tigaiga and T-TE: Teno in Tenerife.

## DISCUSSION

Moulted tail feathers were confirmed to be a valuable non-invasive source of genetic material for both microsatellite genotyping and molecular sexing. Using the universal P8/P2 primer pair for molecular sexing with DNA from *C. bollii* feathers produced a band-size profile characterised by the larger length of Z-intron, consistent with that described for other columbids (Jensen *et al*. 2003; Çakmak *et al*. 2017).

### Genetic diversity and population differentiation

Genetic diversity estimated for *C. bollii* was low. While strong selective constraints may preclude comparisons of microsatellite data across species evolutionary separated, our results show lower diversity in *C. bollii* than observed in other species of Columbiformes (Santiago-Alarcón *et al*. 2006, Pruett *et al*. 2011, Monceau *et al*. 2013), although genetic diversity was similar to that in *C. janthina janthina*, endemic to islands in East Asia (Ando *et al*. 2011). However, the presence of null alleles may have underestimated our results; although the mean frequency of null alleles was relatively low (< 0.1), the average heterozygosity increased after correcting the genotype data for false homozygotes. In addition, all islands/populations and loci were in HWE using the corrected data set. These results suggest that null alleles may be the main cause of heterozygote deficiencies and departures from HWE. Their potential bias should, therefore, be evaluated in population genetics despite their relatively low frequency. However, other factors such as the Wahlund effect (unresolved population substructure), inbreeding or selection near, or at, microsatellite loci might have similar effects on the heterozygosity estimation, so these factors should not be ruled out as contributing to departures from HWE (Dakin & Avise 2004).

Genetic variability within *C. bollii* populations was very similar in all sites (see Tables 1 and S1), suggesting that the populations share a common origin through a metapopulation structure. AMOVA results also support this. However, the global F_ST_ and pairwise genetic differentiation values did reveal significant genetic structuring in the entire sample and between most islands and populations. Although F_ST_^{ENA}^ indicated a higher level of genetic differentiation than F_ST_ and D_est_, the differentiation patterns inferred from these three estimators were largely consistent: La Palma and El Hierro populations were genetically more similar to each other than to the La Gomera and Tenerife populations. Moreover, AMOVA showed that although most of the genetic variation was explained by within-population variance, weak but significant genetic structure was explained by variation among islands.

These results imply that *C. bollii* populations do not comprise a single panmictic unit over their whole range, but rather of at least three insular groups: La Palma-El Hierro, La Gomera and Tenerife. These results could reflect that *C. bollii* populations once formed a genetically homogeneous population that became differentiated due to physical barriers (open sea and highly fragmented habitat), or, alternatively, to a single colonisation event occurring on an only island (perhaps Tenerife), followed by stepwise dispersal to the neighbouring islands.

The genetic divergence associated with morphological adaptations within and among island populations (eg. Milá *et al*. 2010) has not been explored in *C. bollii*, but some evidence from plumage characteristics suggests that this should be studied in detail (P. Marrero, pers. obs.). Interestingly, phenotypic differentiation as response to insularity has been described in the Azores woodpigeon, *C. palumbus azorica* (Andrade *et al*. 2020), as well as in some Macaronesian songbirds (eg. Dietzen et al. 2006, Päckert *et al*. 2006, Illera *et al*. 2007, 2018).

*Columba bollii* populations clustered together similarly in the NJ tree, PCA and Bayesian analyses. STRUCTURE and GENECLASS2 produced the same general assignment pattern, even when the microsatellite locus with the highest null allele frequency was excluded from the analysis (see Carlsson 2008). Four genetic groups corresponding to the four islands were identified, although La Palma and El Hierro populations may belong to the same cluster according to results from the corrected data set. The STRUCTURE analysis also revealed fine-scale structuring within Tenerife and La Gomera, which should be assessed through more exhaustive sampling. Nevertheless, some degree of contemporary gene flow between pigeons from different islands was detected by GENECLASS2.

The Mantel test also indicated that genetic differentiation can be only partially explained due to isolation by distance. This supports the idea that factors other than geographical proximity influence gene flow between populations. Genetic structuring of populations was corroborated by the Population Graph analysis. The connectivity network revealed moderate and differential gene flow, confirming that genetic divergence between populations is not only because of the spatial distances. The graphical topology generated by the null allele corrected data set yielded fewer edges, implying that the null alleles lead to overestimating the connectivity between populations, in accordance with the findings of Klütsch *et al*. (2012).

The metapopulation structure suggested for *C. bollii* pigeons implies a complex interaction between spatially separated individuals via dispersal. Migrants could also be able to recolonise depleted and/or new areas, a phenomenon that has probably occurred throughout their evolutionary history. Moreover, we also suggest that this population structure tends to reflect the relationship between the islands’ geological ages and the evolution of pigeon populations. Valente *et al*. (2017) estimated that *C. bollii* colonised during the Pleistocene (2.14 Myr; 95% CI: 1.42-2.91 Myr). Although the eastern islands and La Gomera emerged between 24-12 Ma (Carracedo & Troll 2021), geological evidence from the Mio-Pliocene suggests that around 11.6-3.2 Ma Tenerife was comprised of three separate proto-islands (Roque del Conde, Teno and Anaga) that finally merged into one single island between 1.9-0.2 Ma (Ancochea *et al*. 1990, Thirlwall *et al*. 2000), through volcanic eruption and growth of the successive central Las Cañadas edifices. La Palma and El Hierro emerged within the last 3.5 and 1.1 Ma, respectively (Guillou *et al*. 1996; Carracedo *et al*. 2001). The geological history of the archipelago was dominated by intense volcanic eruptions, formation of new terrain, large-scale landslides and variations in sea level (Carracedo 1994), which led to the creation of barriers, local extinctions and re-colonization processes that influenced the evolution of many species including Bolle’s pigeon. Therefore, the genetic traits of *C. bollii* represent ancestral evolutionary responses to the laurel forest distribution on these islands, highlighting La Gomera and Tenerife (more precisely T-TE area) populations as the oldest lineages, given the greater number of private microsatellite alleles detected.

Several passerine species on these islands show a similar genetic structure pattern - with a close relationship between El Hierro and La Palma populations - to that found in Bolle’s pigeons (Illera *et al*. 2012, 2016, Valente *et al*. 2017). The main explanation for this result could be related to the recent divergence of lineages (see above the geological age of these two islands), combined with differences in evolutionary rates in neutral genetic loci (Marthinsen *et al*. 2008).

Finally, our study provides a first approach to investigate the role that large-bodied frugivores, like pigeons, could have played in the biogeography and evolution of some fleshy-fruited plant species, but also in the spread of forest areas (Jordano & Godoy 2002). Through seed dispersal, large-sized fruits (eg. Lauraceae fruits) are able to reach and colonise new territories, contributing to the spatial contribution of genetic variation of plant populations (Corlett 2017).

### Implications for conservation

This study provides a useful first step in gaining the scientific data required to inform appropriate conservation measures for *C. bollii*. The species is now listed as Least Concern by IUCN (2022) with an increasing population trend, possibly due to the recovery of the laurel forests. However, island-endemic columbid species are particularly vulnerable to ecological disturbances and environment changes, such as habitat loss and fragmentation, extreme climatic events and invasive exotic species (Walker 2007).

From the genetic data obtained here, we conclude that the similar genetic diversity found in the pigeon populations may indicate gene flow among them, but not enough to eradicate such population differences. These genetic differences are likely to reflect both the ancestral origins and ecological relationships that should be preserved. Although our results indicate that the pigeon has long-distance flight ability, which has been recently confirmed by some field observations of *C. bollii* individuals in forestal areas of Gran Canaria (Martín *et al*. 2020), the actual range and frequency of dispersal remain unknown. The same lack of information applies regarding data on population size, reproduction, movement patterns and demographic trends including spatiotemporal variation and local adaptations in *C. bollii*. Therefore, the major aim of any management action on Bolle’s pigeons should begin with careful field studies leading to protect ecological diversity and evolutionary processes across the geographical range of the species, as well as to maintain the natural network of genetic connections between their populations (see Snyder *et al*. 1996, Crandall *et al*. 2000, Theodorou *et al*. 2009).

We especially thank Francisco Marrero and Manuel Nogales, who made important contributions to the fieldwork. In addition to Julia Rodríguez, Juan José Marrero, Celia Marrero, Félix M. Medina, Segundo Lorenzo, Félix Rodríguez, Alfonso Quintero, Francisco Martín, Beneharo Rodríguez, Airam Rodríguez, Alejandro Padrón, David P. Padilla, Nuria Macías and Salvador de la Cruz for help with sample collection, and Lewis Spurgin for logistic support in the lab. Miriam González and Carolina Medina assisted in the pigeon dissections and Pedro Jordano helped us with some data analysis.

Vicente M. Cabrera and Brent Emerson made useful comments on a previous version of the manuscript. Special thanks to Ángel Fernández and Antonio Zamorano for logistic support in the field (Garajonay National Park) and to Paulo Oliveira (Madeira Natural Park) for providing us some carcasess of *C. trocaz* for the sexing study. Fieldwork was carried out under scientific permits issued by the Canary Government (Ref: 2011/1343) and the island Cabildos of Tenerife (FYT 223/11), La Palma (Ref: 2011/1840), La Gomera (Ref: 353361) and El Hierro (Ref: MRR/rsh).

## AUTHOR CONTRIBUTIONS

**Patricia Marrero:** conceptualization (lead); formal analysis (lead); investigation (lead); methodology (lead); visualization (lead); writing – original draft (lead); writing – review & editing (lead). **Rosa Fregel:** methodology (supporting); writing – review & editing (equal). **David S. Richardson:** methodology (supporting); writing – review & editing (equal).

## ETHICAL NOTE

None

## FUNDING

The manuscript was edited by Guido Jones, currently funded by the Cabildo de Tenerife, under the TFinnova Programme supported by MEDI and FDCAN funds. P. Marrero and R. Fregel were supported by postdoctoral grants from the Spanish Education Ministry (EX2009-950) and Fundación Canaria Manuel Morales, respectively.

## Data Availability Statement

All the data are provided as supplementary material.

## Notes

### Competing Interest Statement

The authors have declared no competing interest.

