## Supplementary material for "Genetic structure and differentiation of the endemic Bolle’s Laurel Pigeon (*Columba bollii*) in the Canary Islands": Figure S1.docx

**SUPPORTING INFORMATION**

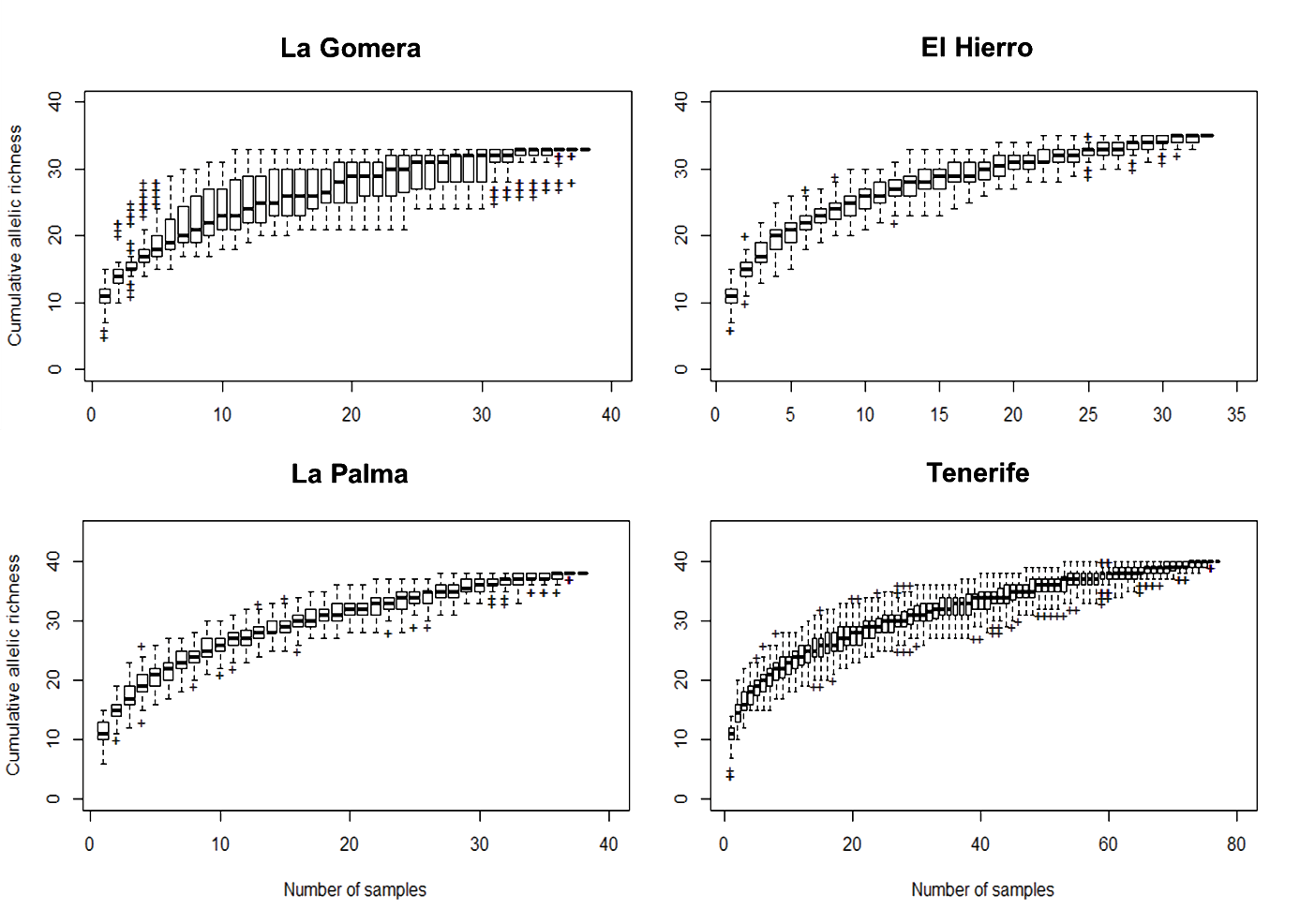

**Figure S1**. Rarefaction curves and confidence intervals of the cumulative allelic richness (mean number of alleles/locus) for feather samples of Bolle's Laurel Pigeon (*Columba bollii*) collected across the Canarian archipelago.
