## Supplementary material for "Genetic structure and differentiation of the endemic Bolle’s Laurel Pigeon (*Columba bollii*) in the Canary Islands": Figure S2.docx

**SUPPORTING INFORMATION**

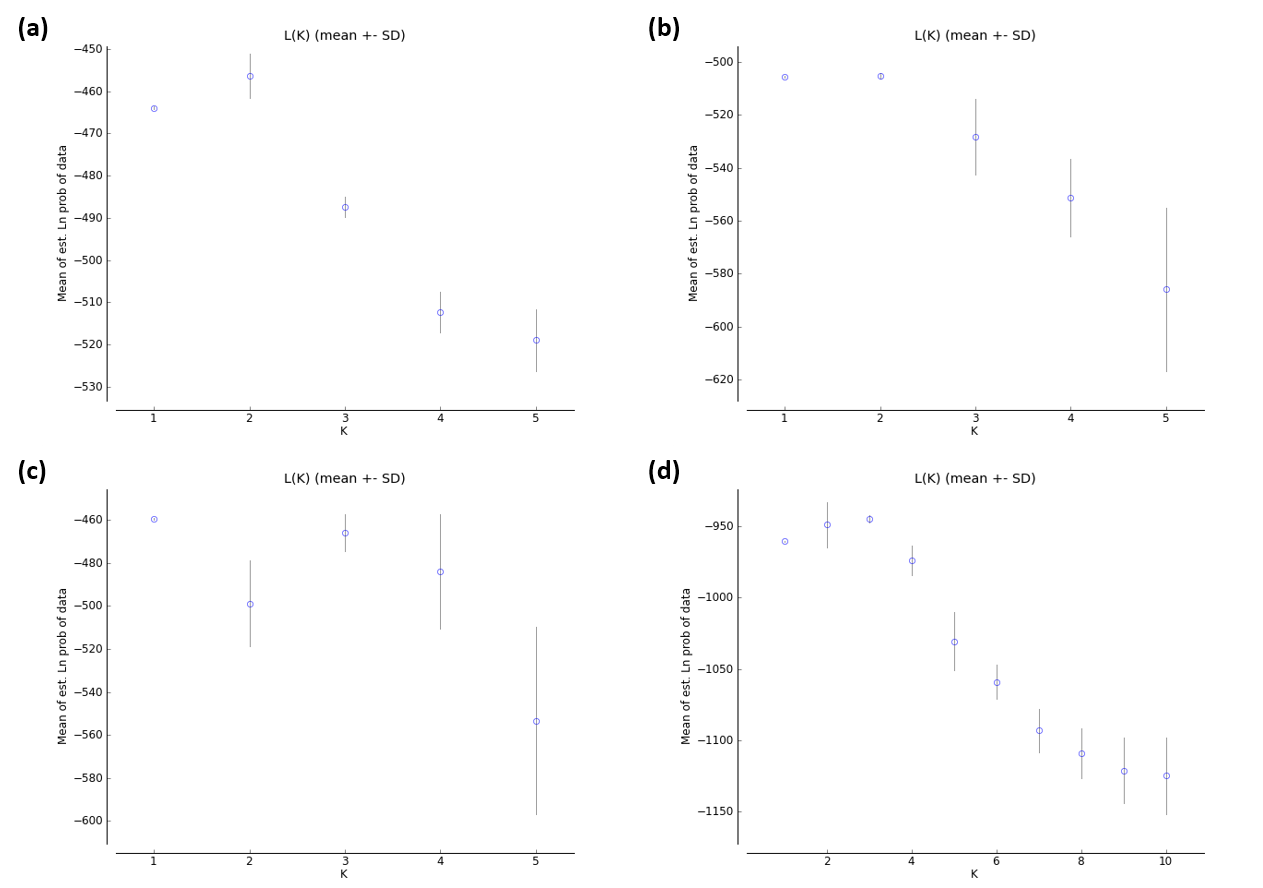

**Figure S2**. Ln values of probability for the assumed number of genetic clusters of Bolle's Laurel Pigeon (*Columba bollii*) populations in (a) El Hierro, (b) La Palma, (c) La Gomera and (d) Tenerife.
