## Supplementary material for "Genetic structure and differentiation of the endemic Bolle’s Laurel Pigeon (*Columba bollii*) in the Canary Islands": Figure S3.docx

**SUPPORTING INFORMATION**

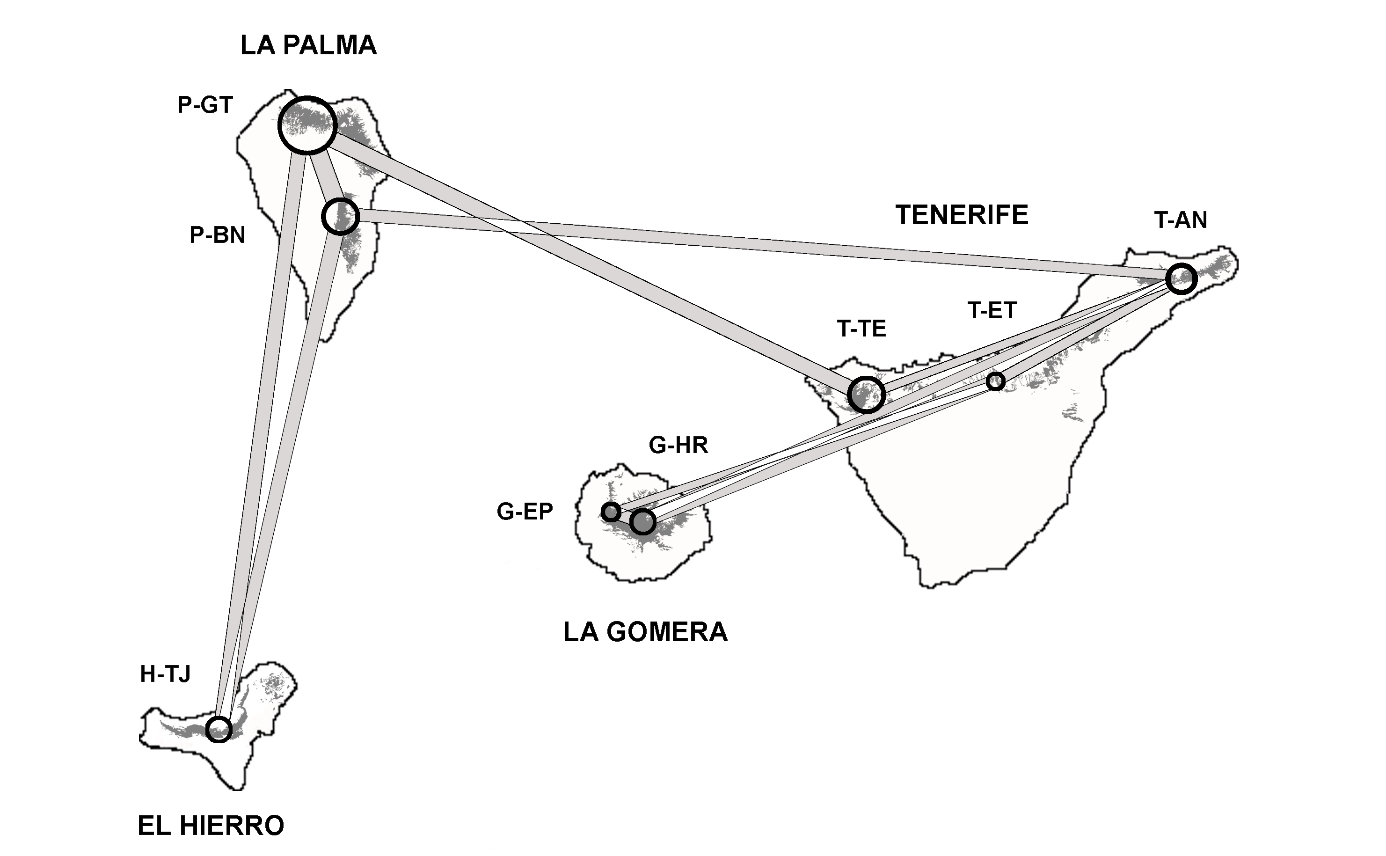

**Figure S3**. Population graph analysis for Bolle's Laurel Pigeon (*Columba bollii*) populations using the corrected data set. Dark grey areas represent the current areas of laurel forests (Del Arco *et al*. 2010) and circles are the central location of sampled populations (size is proportional to within-population allelic richness). The grey edge set indicates the genetic structure conditionally dependent among populations (line width shows the strength of genetic covariance). Sampling localities are indicated as H-JT: Jinama-Tina de las Casillas in El Hierro, P-BN: Barlovento-Niquiomo and P-GT: Garafía-Tinizara in La Palma, G-EP: Epina-Los Pajaritos and G-HR: Hoya del Tión-Los Roques in La Gomera, and T-AN: Anaga, T-ET: La Esperanza-Tigaiga and T-TE: Teno in Tenerife.
