## Supplementary material for "Genetic structure and differentiation of the endemic Bolle’s Laurel Pigeon (*Columba bollii*) in the Canary Islands": Table S1.docx

**SUPPORTING INFORMATION**

**Table S1.** Genetic diversity of ten microsatellite loci among Bolle’s Laurel pigeon (*C. bollii*) populations, *N* = number of samples; *A* = mean number of alleles per locus; *Ar* = mean allelic richness; *Ap* = number of private alleles; *H_O_* = observed heterozygosity; *H_E_* = unbiased expected heterozygosity, *F_IS_* = fixation index, HWE = deviation from Hardy-Weinberg equilibrium after sequential Bonferroni correction (**P* < 0.05, ***P* < 0.01, ****P* < 0.001, n.s. = not significant) and *ȓ_D_* = frequency of null alleles. ^C^ Statistics corrected for null alleles.

| Island | Locus | *A* | *Ar* | *Ap* | *H_O_/* ^C^*H_O_* | *H_E_/* ^C^*H_E_* | *F_IS_/* ^C^*F_IS_* | HWE*/*^C^HWE | *ȓ_D_* |
| --- | --- | --- | --- | --- | --- | --- | --- | --- | --- |
| El Hierro | TG01-077 | 3.00 | 3.00 | 0.00 | 0.515/0.606 | 0.608/0.646 | 0.155/0.062 | n.s./n.s. | 0.056 |
| N = 33 | TG01-124 | 2.00 | 1.85 | 0.00 | 0.043/0.043 | 0.043/0.043 | 0.000/0.000 | n.s./n.s. | 0.000 |
|  | TG01-148 | 3.00 | 2.99 | 0.00 | 0.152/0.515 | 0.338/0.497 | 0.556/-0.036 | **/n.s. | 0.173 |
|  | TG02-078 | 8.00 | 7.54 | 1.00 | 0.194/0.742 | 0.675/0.749 | 0.717/0.010 | ***/n.s. | 0.291 |
|  | TG02-120 | 6.00 | 5.57 | 1.00 | 0.313/0.719 | 0.646/0.727 | 0.520/0.012 | ***/n.s. | 0.208 |
|  | TG03-002 | 3.00 | 2.70 | 1.00 | 0.485/0.485 | 0.438/0.438 | -0.109/-0.109 | n.s./n.s. | 0.000 |
|  | TG04-012 | 5.00 | 4.92 | 0.00 | 0.500/0.656 | 0.542/0.584 | 0.079/-0.125 | n.s./n.s. | 0.067 |
|  | TG07-022 | 2.00 | 2.00 | 0.00 | 0.037/0.037 | 0.037/0.037 | 0.000/0.000 | n.s./n.s. | 0.000 |
|  | TG13-017 | 3.00 | 2.79 | 0.00 | 0.207/0.207 | 0.192/0.192 | -0.080/-0.080 | n.s./n.s. | 0.000 |
|  | TG22-001 | 2.00 | 2.00 | 0.00 | 0.103/0.354 | 0.216/0.379 | 0.525/0.092 | n.s./n.s. | 0.126 |
| La Palma | TG01-077 | 3.00 | 3.00 | 0.00 | 0.432/0.432 | 0.504/0.504 | 0.143/0.143 | n.s./n.s. | 0.054 |
| N = 38 | TG01-124 | 1.00 | 1.00 | 0.00 | 0.030/0.242 | 0.089/0.246 | 0.663/0.015 | n.s./n.s. | 0.109 |
|  | TG01-148 | 3.00 | 2.64 | 0.00 | 0.250/0.444 | 0.372/0.477 | 0.331/0.069 | n.s./n.s. | 0.099 |
|  | TG02-078 | 8.00 | 6.99 | 1.00 | 0.316/0.684 | 0.598/0.696 | 0.475/0.017 | */n.s. | 0.177 |
|  | TG02-120 | 6.00 | 5.49 | 1.00 | 0.711/0.763 | 0.685/0.698 | -0.038/-0.095 | n.s./n.s. | 0.022 |
|  | TG03-002 | 2.00 | 2.00 | 0.00 | 0.324/0.324 | 0.311/0.311 | -0.043/-0.043 | n.s./n.s. | 0.000 |
|  | TG04-012 | 6.00 | 5.37 | 0.00 | 0.368/0.553 | 0.499/0.571 | 0.265/0.033 | n.s./n.s. | 0.090 |
|  | TG07-022 | 3.00 | 2.61 | 0.00 | 0.000/0.000 | 0.000/0.000 | NA | NA | 0.001 |
|  | TG13-017 | 4.00 | 3.43 | 1.00 | 0.143/0.143 | 0.138/0.138 | -0.037/-0.037 | n.s./n.s. | 0.000 |
|  | TG22-001 | 3.00 | 2.97 | 1.00 | 0.121/0.394 | 0.223/0.386 | 0.461/-0.022 | n.s./n.s. | 0.124 |
| La Gomera | TG01-077 | 3.00 | 3.00 | 0.00 | 0.405/0.595 | 0.526/0.599 | 0.231/0.008 | n.s./n.s. | 0.085 |
| N = 38 | TG01-124 | 2.00 | 2.00 | 0.00 | 0.114/0.314 | 0.205/0.354 | 0.447/0-090 | n.s./n.s. | 0.108 |
|  | TG01-148 | 5.00 | 4.15 | 2.00 | 0.250/0.472 | 0.371/0.488 | 0.330/0.033 | ***/n.s. | 0.108 |
|  | TG02-078 | 5.00 | 4.23 | 0.00 | 0.324/0.324 | 0.292/0.292 | -0.111/-0.111 | n.s./n.s. | 0.000 |
|  | TG02-120 | 5.00 | 4.86 | 0.00 | 0.306/0.694 | 0.584/0.687 | 0.480/-0.010 | ***/n.s. | 0.185 |
|  | TG03-002 | 5.00 | 3.82 | 3.00 | 0.316/0.474 | 0.450/0.539 | 0.301/0.123 | ***/n.s. | 0.087 |
|  | TG04-012 | 4.00 | 3.90 | 1.00 | 0.324/0.351 | 0.351/0.372 | 0.077/0.057 | n.s./n.s. | 0.008 |
|  | TG07-022 | 3.00 | 2.90 | 1.00 | 0.176/0.441 | 0.304/0.452 | 0.423/0.025 | */n.s. | 0.131 |
|  | TG13-017 | 2.00 | 1.99 | 0.00 | 0.059/0.235 | 0.112/0.241 | 0.480/0.024 | n.s./n.s. | 0.091 |
|  | TG22-001 | 2.00 | 1.89 | 0.00 | 0.057/0.057 | 0.056/0.056 | -0.015/-0.015 | n.s./n.s. | 0.000 |

**Table S1.** Continued.

|  | Locus | A | Ar | Ap | H_O_/ ^C^H_O_ | H_E_/ ^C^H_E_ | F_IS_/ ^C^F_IS_ | HWE/^C^HWE | ȓ_D_ |
| --- | --- | --- | --- | --- | --- | --- | --- | --- | --- |
| Tenerife | TG01-077 | 4.00 | 3.29 | 1.00 | 0.312/0.571 | 0.491/0.592 | 0.367/0.035 | **/n.s. | 0.133 |
| N = 77 | TG01-124 | 4.00 | 2.98 | 1.00 | 0.069/0.264 | 0.133/0.273 | 0.479/0.034 | */n.s. | 0.095 |
|  | TG01-148 | 4.00 | 3.43 | 2.00 | 0.438/0.630 | 0.521/0.583 | 0.160/-0.082 | ***/* | 0.099 |
|  | TG02-078 | 4.00 | 3.89 | 0.00 | 0.261/0.580 | 0.460/0.582 | 0.435/0.003 | ***/n.s. | 0.154 |
|  | TG02-120 | 7.00 | 5.13 | 2.00 | 0.421/0.592 | 0.532/0.599 | 0.209/0.012 | n.s./n.s. | 0.084 |
|  | TG03-002 | 2.00 | 2.00 | 0.00 | 0.312/0.442 | 0.387/0.458 | 0.196/0.037 | n.s./n.s. | 0.060 |
|  | TG04-012 | 7.00 | 4.22 | 1.00 | 0.467/0.467 | 0.464/0.464 | -0.007/-0.007 | n.s./n.s. | 0.000 |
|  | TG07-022 | 3.00 | 2.68 | 1.00 | 0.074/0.309 | 0.152/0.308 | 0.519/-0.002 | **/n.s. | 0.116 |
|  | TG13-017 | 3.00 | 2.48 | 0.00 | 0.042/0.292 | 0.119/0.291 | 0.653/-0.003 | ***/n.s. | 0.124 |
|  | TG22-001 | 4.00 | 2.82 | 2.00 | 0.097/0.264 | 0.145/0.263 | 0.329/-0.002 | n.s./n.s. | 0.079 |
| Population |  |  |  |  |  |  |  |  |  |
| H-JT | TG01-077 | 3.00 | 3.00 | 0.00 | 0.515/0.606 | 0.608/0.646 | 0.155/0.062 | n.s./n.s. | 0.056 |
| N = 33 | TG01-124 | 2.00 | 2.00 | 0.00 | 0.043/0.043 | 0.043/0.043 | 0.000/0.000 | n.s./n.s. | 0.000 |
|  | TG01-148 | 3.00 | 2.99 | 0.00 | 0.152/0.515 | 0.338/0.497 | 0.556/-0.036 | **/n.s. | 0.173 |
|  | TG02-078 | 8.00 | 7.54 | 1.00 | 0.194/0.742 | 0.675/0.749 | 0.717/0.010 | ***/n.s. | 0.291 |
|  | TG02-120 | 6.00 | 5.57 | 1.00 | 0.313/0.719 | 0.646/0.727 | 0.520/0.012 | ***/n.s. | 0.208 |
|  | TG03-002 | 3.00 | 2.70 | 1.00 | 0.485/0.485 | 0.438/0.438 | -0.109/-0.109 | n.s./n.s. | 0.000 |
|  | TG04-012 | 5.00 | 4.92 | 0.00 | 0.500/0.656 | 0.542/0.584 | 0.079/-0.125 | n.s./n.s. | 0.067 |
|  | TG07-022 | 2.00 | 1.85 | 0.00 | 0.037/0.037 | 0.037/0.037 | 0.000/0.000 | n.s./n.s. | 0.000 |
|  | TG13-017 | 3.00 | 2.79 | 0.00 | 0.207/0.207 | 0.192/0.192 | -0.080/-0.080 | n.s./n.s. | 0.000 |
|  | TG22-001 | 2.00 | 2.00 | 0.00 | 0.103/0.345 | 0.216/0.379 | 0.525/0.092 | n.s./n.s. | 0.126 |
| P-BN | TG01-077 | 3.00 | 3.00 | 0.00 | 0.450/0.650 | 0.545/0.614 | 0.178/-0.060 | n.s./n.s. | 0.074 |
| N = 21 | TG01-124 | 1.00 | 1.00 | 0.00 | 0.000/0.000 | 0.000/0.000 | NA | NA | 0.001 |
|  | TG01-148 | 2.00 | 2.00 | 0.00 | 0.250/0.450 | 0.358/0.472 | 0.307/0.047 | n.s./n.s. | 0.087 |
|  | TG02-078 | 7.00 | 5.58 | 0.00 | 0.190/0.667 | 0.534/0.668 | 0.649/0.002 | ***/n.s. | 0.225 |
|  | TG02-120 | 5.00 | 4.57 | 0.00 | 0.762/0.762 | 0.722/0.722 | -0.056/-0.056 | n.s./n.s. | 0.000 |
|  | TG03-002 | 2.00 | 2.00 | 0.00 | 0.381/0.381 | 0.372/0.372 | -0.026/-0.026 | n.s./n.s. | 0.000 |
|  | TG04-012 | 5.00 | 4.34 | 0.00 | 0.429/0.429 | 0.410/0.410 | -0.047/-0.047 | n.s./n.s. | 0.000 |
|  | TG07-022 | 1.00 | 1.00 | 0.00 | 0.000/0.000 | 0.000/0.000 | NA | NA | 0.001 |
|  | TG13-017 | 3.00 | 2.44 | 0.00 | 0.111/0.111 | 0.110/0.110 | -0.015/-0.015 | n.s./n.s. | 0.000 |
|  | TG22-001 | 2.00 | 1.99 | 0.00 | 0.105/0.316 | 0.193/0.326 | 0.463/0.031 | n.s./n.s. | 0.107 |
| P-GT | TG01-077 | 3.00 | 2.99 | 0.00 | 0.412/0.412 | 0.465/0.465 | 0.118/0.118 | n.s./n.s. | 0.010 |
| N = 17 | TG01-124 | 3.00 | 2.93 | 0.00 | 0.071/0.357 | 0.204/0.373 | 0.658/0.044 | n.s./n.s. | 0.154 |
|  | TG01-148 | 3.00 | 2.81 | 0.00 | 0.250/0.500 | 0.401/0.534 | 0.385/0.066 | n.s./n.s. | 0.113 |
|  | TG02-078 | 6.00 | 5.29 | 1­.00 | 0.471/0.706 | 0.663/0.731 | 0.297/0.035 | n.s./n.s. | 0.104 |
|  | TG02-120 | 5.00 | 4.71 | 1.00 | 0.647/0.765 | 0.624/0.656 | -0.038/-0.172 | n.s./n.s. | 0.045 |
|  | TG03-002 | 2.00 | 2.00 | 0.00 | 0.250/0.250 | 0.226/0.226 | -0.111/-0.111 | n.s./n.s. | 0.000 |
|  | TG04-012 | 5.00 | 4.52 | 0.00 | 0.294/0.647 | 0.592/0.692 | 0.511/0.066 | ***/n.s. | 0.181 |
|  | TG07-022 | 1.00 | 1.00 | 0.00 | 0.000/0.000 | 0.000/0.000 | NA | NA | 0.001 |
|  | TG13-017 | 4.00 | 3.29 | 0.00 | 0.176/0.176 | 0.171/0.171 | -0.032/-0.032 | n.s./n.s. | 0.000 |
|  | TG22-001 | 3.00 | 2.93 | 1.00 | 0.143/0.429 | 0.262/0.426 | 0.464/-0.006 | n.s./n.s. | 0.122 |

**Table S1.** Continued.

| Population | Locus | A | Ar | Ap | H_O_/ ^C^H_O_ | H_E_/ ^C^H_E_ | F_IS_/ ^C^F_IS_ | HWE/^C^HWE | ȓ_D_ |
| --- | --- | --- | --- | --- | --- | --- | --- | --- | --- |
| G-EP | TG01-077 | 3.00 | 2.99 | 0.00 | 0.435/0.609 | 0.534/0.596 | 0.190/-0.022 | n.s./n.s. | 0.071 |
| N = 23 | TG01-124 | 2.00 | 1.59 | 0.00 | 0.045/0.045 | 0.045/0.045 | 0.000/0.000 | n.s./n.s. | 0.000 |
|  | TG01-148 | 3.00 | 2.84 | 0.00 | 0.091/0.500 | 0.317/0.496 | 0.718/-0.009 | */n.s. | 0.203 |
|  | TG02-078 | 5.00 | 4.11 | 0.00 | 0.455/0.455 | 0.393/0.393 | -0.160/-0.160 | n.s./n.s. | 0.000 |
|  | TG02-120 | 5.00 | 4.65 | 0.00 | 0.227/0.682 | 0.540/0.667 | 0.585/-0.023 | **/n.s. | 0.209 |
|  | TG03-002 | 2.00 | 2.00 | 0.00 | 0.174/0.478 | 0.394/0.544 | 0.564/0.123 | n.s./n.s. | 0.167 |
|  | TG04-012 | 4.00 | 3.38 | 0.00 | 0.348/0.391 | 0.370/0.404 | 0.061/0.032 | n.s./n.s. | 0.015 |
|  | TG07-022 | 2.00 | 2.00 | 0.00 | 0.238/0.286 | 0.285/0.324 | 0.167/0.121 | n.s./n.s. | 0.043 |
|  | TG13-017 | 2.00 | 1.96 | 0.00 | 0.050/0.300 | 0.142/0.309 | 0.655/0.030 | n.s./n.s. | 0.127 |
|  | TG22-001 | 2.00 | 1.62 | 0.00 | 0.048/0.048 | 0.048/0.048 | 0.000/0.000 | n.s./n.s. | 0.000 |
| G-HR | TG01-077 | 3.00 | 2.93 | 0.00 | 0.357/0.571 | 0.521/0.619 | 0.323/0.080 | n.s./n.s. | 0.095 |
| N = 15 | TG01-124 | 2.00 | 2.00 | 0.00 | 0.231/0.462 | 0.409/0.526 | 0.446/0.127 | n.s./n.s. | 0.128 |
|  | TG01-148 | 5.00 | 4.79 | 3.00 | 0.571/0.500 | 0.508/0.458 | -0.130/-0.096 | ***/n.s. | 0.000 |
|  | TG02-078 | 2.00 | 1.99 | 0.00 | 0.133/0.133 | 0.129/0.129 | -0.037/-0.037 | n.s./n.s. | 0.000 |
|  | TG02-120 | 4.00 | 4.00 | 0.00 | 0.429/0.714 | 0.653/0.725 | 0.353/0.015 | n.s./n.s. | 0.144 |
|  | TG03-002 | 5.00 | 4.60 | 3.00 | 0.533/0.533 | 0.543/0.543 | 0.018/0.018 | ***/n.s. | 0.000 |
|  | TG04-012 | 4.00 | 3.92 | 0.00 | 0.286/0.286 | 0.325/0.325 | 0.126/0.126 | n.s./n.s. | 0.000 |
|  | TG07-022 | 3.00 | 3.00 | 1.00 | 0.077/0.538 | 0.342/0.532 | 0.782/-0.012 | n.s./n.s. | 0.224 |
|  | TG13-017 | 2.00 | 1.93 | 0.00 | 0.071/0.071 | 0.071/0.071 | 0.000/0.000 | n.s./n.s. | 0.000 |
|  | TG22-001 | 2.00 | 1.93 | 0.00 | 0.071/0.071 | 0.071/0.071 | 0.000/0.000 | n.s./n.s. | 0.000 |
| T-AN | TG01-077 | 3.00 | 2.93 | 0.00 | 0.162/0.649 | 0.506/0.647 | 0.679/-0.002 | **/n.s. | 0.237 |
| N = 37 | TG01-124 | 3.00 | 2.25 | 0.00 | 0.088/0.265 | 0.140/0.266 | 0.373/0.007 | n.s./n.s. | 0.084 |
|  | TG01-148 | 4.00 | 3.46 | 1.00 | 0.371/0.686 | 0.566/0.649 | 0.381/-0.058 | ***/n.s. | 0.154 |
|  | TG02-078 | 4.00 | 3.46 | 0.00 | 0.114/0.600 | 0.421/0.591 | 0.730/-0.015 | ***/n.s. | 0.237 |
|  | TG02-120 | 7.00 | 4.66 | 1.00 | 0.444/0.583 | 0.540/0.595 | 0.171/0.019 | n.s./n.s. | 0.069 |
|  | TG03-002 | 2.00 | 2.00 | 0.00 | 0.243/0.405 | 0.359/0.445 | 0.320/0.091 | n.s./n.s. | 0.096 |
|  | TG04-012 | 6.00 | 4.05 | 0.00 | 0.541/0.568 | 0.572/0.589 | 0.067/0.036 | n.s./n.s. | 0.013 |
|  | TG07-022 | 2.00 | 1.83 | 1.00 | 0.033/0.267 | 0.097/0.267 | 0.659/0.002 | n.s./n.s. | 0.109 |
|  | TG13-017 | 2.00 | 1.86 | 0.00 | 0.000/0.324 | 0.112/0.324 | 1.000/0.000 | */n.s. | 0.160 |
|  | TG22-001 | 3.00 | 2.25 | 0.00 | 0.054/0.324 | 0.153/0.327 | 0.649/0.008 | */n.s. | 0.134 |
| T-ET | TG01-077 | 3.00 | 2.91 | 0.00 | 0.421/0.421 | 0.421/0.421 | 0.013/0.000 | n.s./n.s. | 0.000 |
| N = 19 | TG01-124 | 2.00 | 1.68 | 0.00 | 0.053/0.053 | 0.053/0.053 | 0.000/0.000 | n.s./n.s. | 0.000 |
|  | TG01-148 | 2.00 | 2.00 | 0.00 | 0.632/0.632 | 0.478/0.478 | -0.365/-0.333 | n.s./n.s. | 0.000 |
|  | TG02-078 | 4.00 | 3.91 | 0.00 | 0.444/0.611 | 0.548/0.614 | 0.207/0.005 | n.s./n.s. | 0.077 |
|  | TG02-120 | 4.00 | 3.66 | 0.00 | 0.368/0.632 | 0.539/0.634 | 0.335/0.005 | n.s./n.s. | 0.116 |
|  | TG03-002 | 2.00 | 2.00 | 0.00 | 0.579/0.632 | 0.462/0.496 | -0.229/-0.282 | n.s./n.s. | 0.000 |
|  | TG04-012 | 3.00 | 2.68 | 0.00 | 0.263/0.263 | 0.240/0.240 | -0.101/-0.098 | n.s./n.s. | 0.000 |
|  | TG07-022 | 2.00 | 2.00 | 0.00 | 0.056/0.444 | 0.246/0.446 | 0.780/0.004 | n.s./n.s. | 0.186 |
|  | TG13-017 | 3.00 | 2.59 | 0.00 | 0.053/0.316 | 0.152/0.324 | 0.661/0.027 | n.s./n.s. | 0.137 |
|  | TG22-001 | 3.00 | 2.44 | 1.00 | 0.111/0.111 | 0.110/0.110 | -0.014/-0.015 | n.s./n.s. | 0.000 |

**Table S1.** Continued.

| Population | Locus | A | Ar | Ap | H_O_/ ^C^H_O_ | H_E_/ ^C^H_E_ | F_IS_/ ^C^F_IS_ | HWE/^C^HWE | ȓ_D_ |
| --- | --- | --- | --- | --- | --- | --- | --- | --- | --- |
| T-TE | TG01-077 | 4.00 | 3.61 | 1.00 | 0.476/0.524 | 0.539/0.569 | 0.119/0.081 | n.s./n.s. | 0.046 |
| N = 21 | TG01-124 | 4.00 | 3.28 | 1.00 | 0.053/0.368 | 0.201/0.401 | 0.743/0.084 | ***/n.s. | 0.148 |
|  | TG01-148 | 3.00 | 2.91 | 0.00 | 0.368/0.579 | 0.494/0.568 | 0.259/-0.021 | n.s./n.s. | 0.112 |
|  | TG02-078 | 4.00 | 3.81 | 0.00 | 0.375/0.500 | 0.464/0.510 | 0.196/0.020 | n.s./n.s. | 0.066 |
|  | TG02-120 | 4.00 | 3.62 | 0.00 | 0.429/0.524 | 0.519/0.580 | 0.178/0.098 | n.s./n.s. | 0.059 |
|  | TG03-002 | 2.00 | 2.00 | 0.00 | 0.190/0.476 | 0.372/0.511 | 0.494/0.070 | n.s./n.s. | 0.143 |
|  | TG04-012 | 4.00 | 3.59 | 1.00 | 0.526/0.526 | 0.431/0.431 | -0.229/-0.229 | n.s./n.s. | 0.000 |
|  | TG07-022 | 2.00 | 1.96 | 0.00 | 0.150/0.150 | 0.142/0.142 | -0.056/-0.056 | n.s./n.s. | 0.000 |
|  | TG13-017 | 2.00 | 1.91 | 0.00 | 0.105/0.105 | 0.102/0.102 | -0.029/-0.029 | n.s./n.s. | 0.000 |
|  | TG22-001 | 3.00 | 2.72 | 0.00 | 0.176/0.176 | 0.169/0.169 | -0.043/-0.043 | n.s./n.s. | 0.000 |

Sampling localities are indicated as H-JT: Jinama-Tina de las Casillas in El Hierro, P-BN: Barlovento-Niquiomo and P-GT: Garafía-Tinizara in La Palma, G-EP: Epina-Los Pajaritos and G-HR: Hoya del Tión-Los Roques in La Gomera, and T-AN: Anaga, T-ET: La Esperanza-Tigaiga and T-TE: Teno in Tenerife.
