## Supplementary material for "Genetic structure and differentiation of the endemic Bolle’s Laurel Pigeon (*Columba bollii*) in the Canary Islands": Table S2.docx

**SUPPORTING INFORMATION**

**Table S2.** Pairwise *F_ST_* (below diagonal) and pairwise *F_ST_^{ENA}^* with 95% confidence intervals (above diagonal) between all populations of Bolle’s Laurel pigeon on the Canary Islands. Statistical significance after sequential Bonferroni correction is indicated (**P* < 0.05, ***P* <0.01, n.s. = not significant).

|  | H-JT | P-BN | P-GT | G-EP | G-HR | T-AN | T-ET | T-TE |
| --- | --- | --- | --- | --- | --- | --- | --- | --- |
| H-JT |  | -0.002 (-0.009-0.006)^n.s.^ | 0.013 (-0.001-0.031)^n.s.^ | 0.029 (0.0003-0.067)** | 0.072 (0.009-0.156)** | 0.024 (0.009-0.041)** | 0.041 (0.019-0.067)** | 0.019 (0.006-0.035)* |
| P-BN | 0.010 ^n.s.^ |  | 0.013 (-0.009-0.038)^n.s.^ | 0.022 (-0.0003-0.050)** | 0.054 (-0.006-0.138)** | 0.025 (0.0005-0.047)* | 0.024 (0.005-0.054)* | 0.012 (-0.008-0.035)^n.s.^ |
| P-GT | 0.016 ^n.s.^ | 0.019 ^n.s.^ |  | 0.045 (0.011-0.081)** | 0.069 (0.013-0.134)** | 0.020 (0.004-0.035)* | 0.049 (0.005-0.107)** | 0.016 (-0.009-0.048)^n.s.^ |
| G-EP | 0.017 ^n.s.^ | 0.019 ^n.s.^ | 0.027 ^n.s.^ |  | ^.^0.015 (-0.014-0.072)^n.s.^ | 0.023 (0.001-0.046)** | 0.010 (-0.008-0.031)^n.s.^ | 0.006 (-0.010-0.028)^n.s.^ |
| G-HR | 0.037 ^n.s.^ | 0.031 ^n.s.^ | 0.044 ^n.s.^ | 0.021^n.s^ |  | 0.047 (0.017-0.082)** | 0.039 (-0.005-0.097)* | 0.019 (-0.005-0.053)^n.s.^ |
| T-AN | 0.022 ^n.s.^ | 0.021 ^n.s.^ | 0.022 ^n.s.^ | 0.021 ^n.s.^ | 0.030 ^n.s.^ |  | 0.017 (-0.002-0.041)* | -0.002 (-0.009-0.006)^n.s.^ |
| T-ET | 0.025 ^n.s.^ | 0.021 ^n.s.^ | 0.030 ^n.s.^ | 0.017 ^n.s.^ | 0.030 ^n.s.^ | 0.015 ^n.s.^ |  | 0.006 (-0.007-0.027)** |
| T-TE | 0.015 ^n.s.^ | 0.015 ^n.s.^ | 0.020 ^n.s.^ | 0.012 ^n.s.^ | 0.023 ^n.s.^ | 0.006 ^n.s.^ | 0.010 ^n.s.^ |  |

Sampling localities are indicated as H-JT: Jinama-Tina de las Casillas in El Hierro, P-BN: Barlovento-Niquiomo and P-GT: Garafía-Tinizara in La Palma, G-EP: Epina-Los Pajaritos and G-HR: Hoya del Tión-Los Roques in La Gomera, and T-AN: Anaga, T-ET: La Esperanza-Tigaiga and T-TE: Teno in Tenerife.
